## Supplementary Figures for "Bone morphogenetic protein (BMP) signaling determines neuroblastoma cell fate and sensitivity to retinoic acid"

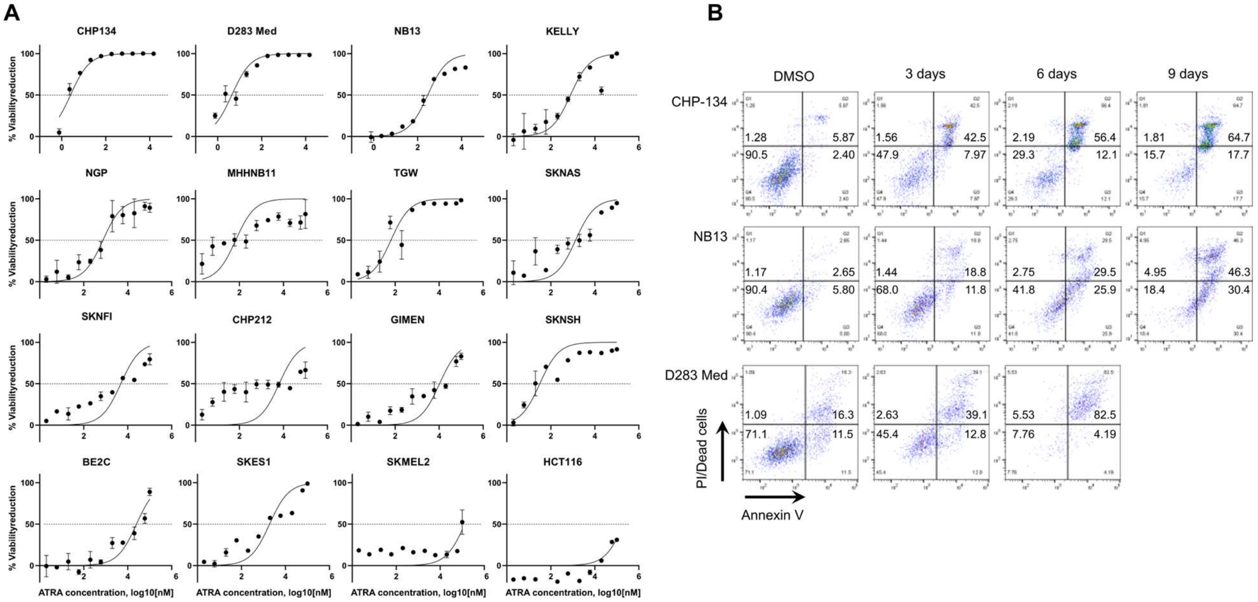


C

| Index | Name | P-value | Adjusted p-value | Odds Ratio | Combined score |
|---|---|---|---|---|---|
| 1 | ALK2 pathway | 1.04E-06 | 0.000149 | 43.35 | 597.18 |
| 2 | BMP signaling and regulation | 1.76E-06 | 0.000205 | 37.16 | 492.41 |
| 3 | Signaling by bone morphogenetic protein (BMP) | 1.65E-07 | 5.44E-05 | 22.87 | 357.11 |
| 4 | BMP signaling pathway in stem cells | 6.35E-06 | 0.000495 | 26.01 | 311.2 |
| 5 | p27 phosphorylation regulation during cell cycle progression | 8.17E-05 | 0.002604 | 23.06 | 217.01 |
| 6 | G2 phase pathway | 0.003502 | 0.03458 | 34.41 | 194.58 |
| 7 | TGF-beta signaling in development | 7.78E-06 | 0.000545 | 15.64 | 183.96 |
| 8 | Cell cycle: G1/S checkpoint | 1.23E-05 | 0.000617 | 14.21 | 160.68 |
| 9 | Rb tumor suppressor/checkpoint signaling in response to DNA damage | 0.000199 | 0.004303 | 17.29 | 147.36 |
| 10 | Transfer RNA aminoacylation | 2.28E-06 | 0.000228 | 11.02 | 143.2 |

D

Depletion (negative selection)

| Gene | negative rank | negative p-value |
|---|---|---|
| *FKBP1A* | 1 | 2.59E-07 |
| *SMURF2* | 20 | 2.04E-04 |
| *SMAD7* | 171 | 4.99E-03 |
| *SMAD6* | 304 | 1.30E-02 |

Enrichment (positive selection)

| Gene | positive rank | positive p-value |
|---|---|---|
| *BMPR2* | 81 | 2.28E-03 |
| *RARA* | 91 | 2.76E-03 |
| *ACVR1* | 178 | 6.62E-03 |
| *SMAD4* | 245 | 0.01 |

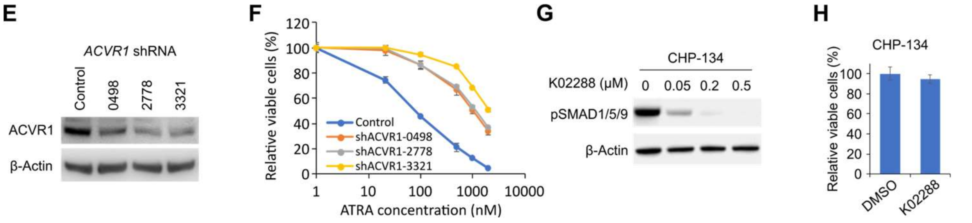


### Supplementary Figure 1

A. Dose response curve of ATRA in 12 neuroblastoma cell lines and 4 non-neuroblastoma cell lines. The data were retrieved from Lee et al (PMID 37957169) (Lee, Wright et al. 2023), except CHP-134, D283 Med, and NB13. The curve fit was performed using GraphPad Prism 9 (nonlinear regression, log(agonist) vs. normalized response). Data represent the mean ± SEM.
B. Flow cytometry results showing apoptosis in CHP-134, NB13, and D283 Med cell lines treated with RA (2μM). Apoptotic cells were stained positive with annexin V and dead cells were stained positive with PI. Representative results of three independent replicates are shown.
C. Table of the top pathways identified from the genome-wide CRISPR knockout screen in CHP-134 cell line, as reported by Enrichr (https://maayanlab.cloud/Enrichr/). Top 1% of

genes with lowest positive RRA score and top 1% of genes with lowest negative RRA score (191 + 191 genes) were used as input.

D. CRISPR knockout screen in CHP-134 cell line treated with 2 μM ATRA for 3 days. BMP-related genes and RARA are shown. Genes are ranked by RRA score (rank #1 has the lowest RRA score).
E. Western blots showing ACVR1 expression in CHP-134 cells following lentiviral shRNA knockdown. Control is a control shRNA with no specific target in human genome; 0498, 2778, and 3321 are three independent shRNAs targeting ACVR1. β-Actin was used as a loading control Representative results of 3 independent replicates are shown.
F. Cell viability in CHP-134 cells with ACVR1 knockdown. Viability was measured with MTS assay and normalized to DMSO treated cells.
G. Western blots showing expression of phosphorylated SMAD1/5/9 in CHP-134 cells treated with K02288 for three days. β-Actin was used as a loading control Representative results of 3 independent replicates are shown.
H. Viability of DMSO and 0.5 μM K02288 treated CHP-134 cells. Cells were treated for 3 days.

For panel F and H, data represent the mean ± SD of 3 independent replicates.

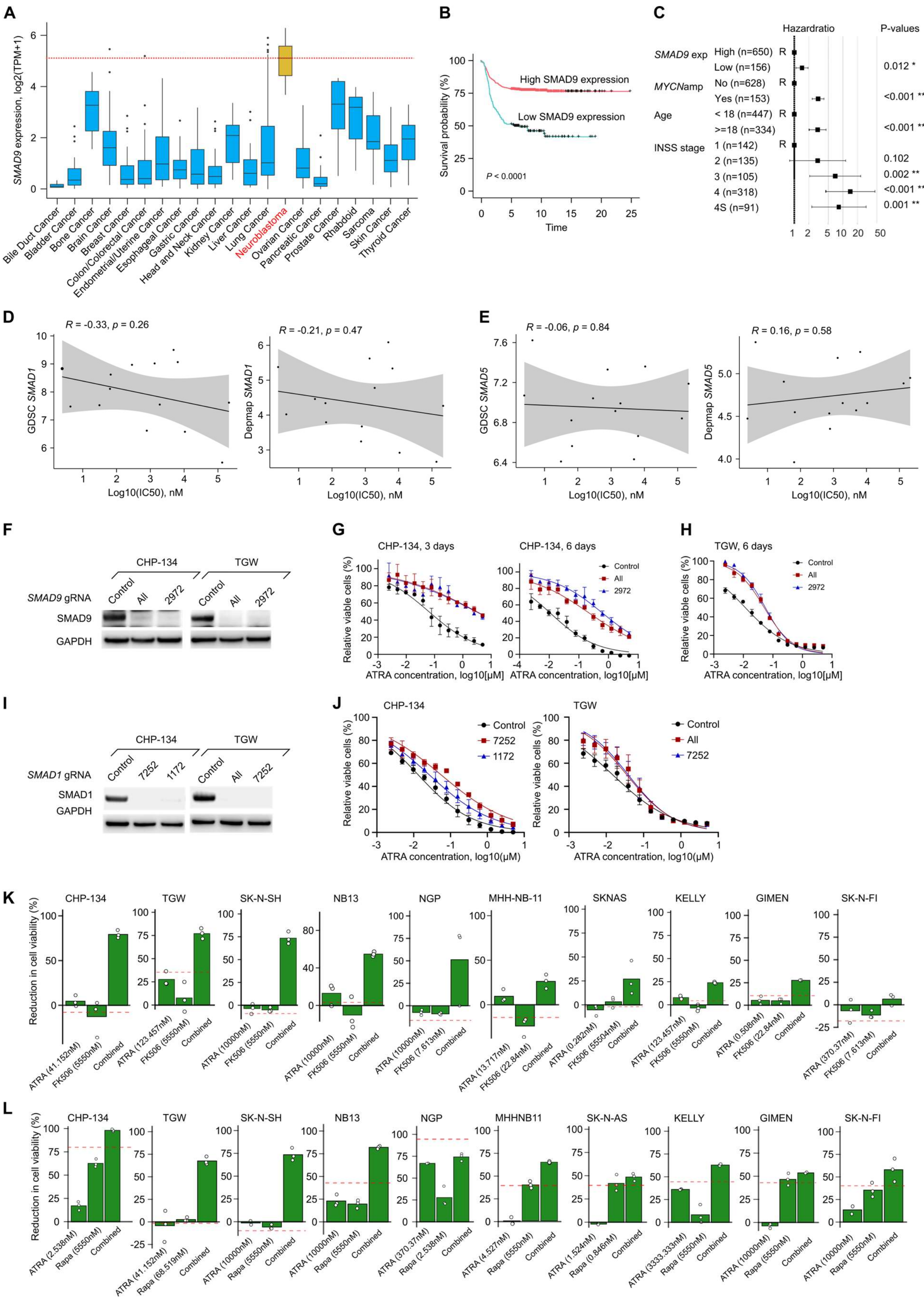
A
SMAD9 expression, log2(TPM+1)
Bile Duct Cancer
Bladder Cancer
Bone Cancer
Brain Cancer
Breast Cancer
Colon/Colorectal Cancer
Endometrial/Uterine Cancer
Esophageal Cancer
Gastric Cancer
Head and Neck Cancer
Kidney Cancer
Liver Cancer
Lung Cancer
Neuroblastoma
Ovarian Cancer
Pancreatic Cancer
Prostate Cancer
Rhabdoid
Sarcoma
Skin Cancer
Thyroid Cancer
B
Survival probability (%)
High SMAD9 expression
Low SMAD9 expression
P < 0.0001
Time
C
Hazardratio
P-values
SMAD9 exp
High (n=650) R
Low (n=156)
0.012 *
MYCNamp
No (n=628) R
Yes (n=153)
<0.001 ***
Age
< 18 (n=447) R
>=18 (n=334)
<0.001 ***
INSS stage
1 (n=142) R
2 (n=135)
0.102
3 (n=105)
0.002 **
4 (n=318)
<0.001 ***
4S (n=91)
0.001 **
D
R = -0.33, p = 0.26
GDSC SMAD1
Log10(IC50), nM
R = -0.21, p = 0.47
Depmap SMAD1
Log10(IC50), nM
E
R = -0.06, p = 0.84
GDSC SMAD5
Log10(IC50), nM
R = 0.16, p = 0.58
Depmap SMAD5
Log10(IC50), nM
F
CHP-134
TGW
SMAD9 gRNA
Control
All
2972
SMAD9
GAPDH
G
CHP-134, 3 days
CHP-134, 6 days
Relative viable cells (%)
Control
All
2972
ATRA concentration, log10[μM]
H
TGW, 6 days
Relative viable cells (%)
Control
All
2972
ATRA concentration, log10[μM]
I
CHP-134
TGW
SMAD1 gRNA
Control
7252
1172
All
SMAD1
GAPDH
J
CHP-134
TGW
Relative viable cells (%)
Control
7252
1172
All
ATRA concentration, log10(μM)
K
Reduction in cell viability (%)
CHP-134
ATRA (41.152nM)
FK506 (5550nM)
Combined
TGW
ATRA (123.457nM)
FK506 (5550nM)
Combined
SK-N-SH
ATRA (10000nM)
FK506 (5550nM)
Combined
NB13
ATRA (10000nM)
FK506 (5550nM)
Combined
NGP
ATRA (10000nM)
FK506 (7.613nM)
Combined
MHH-NB-11
ATRA (13.717nM)
FK506 (22.84nM)
Combined
SKNAS
ATRA (0.282nM)
FK506 (55504nM)
Combined
KELLY
ATRA (123.457nM)
FK506 (5550nM)
Combined
GIMEN
ATRA (0.508nM)
FK506 (22.84nM)
Combined
SK-N-FI
ATRA (370.37nM)
FK506 (7.613nM)
Combined
L
Reduction in cell viability (%)
CHP-134
ATRA (2.538nM)
Rapa (5550nM)
Combined
TGW
ATRA (41.152nM)
Rapa (68.519nM)
Combined
SK-N-SH
ATRA (10000nM)
Rapa (5550nM)
Combined
NB13
ATRA (10000nM)
Rapa (5550nM)
Combined
NGP
ATRA (370.37nM)
Rapa (2.538nM)
Combined
MHHNB11
ATRA (4.527nM)
Rapa (5550nM)
Combined
SK-N-AS
ATRA (1.524nM)
Rapa (0.846nM)
Combined
KELLY
ATRA (3333.333nM)
Rapa (5550nM)
Combined
GIMEN
ATRA (10000nM)
Rapa (5550nM)
Combined
SK-N-FI
ATRA (10000nM)
Rapa (5550nM)
Combined

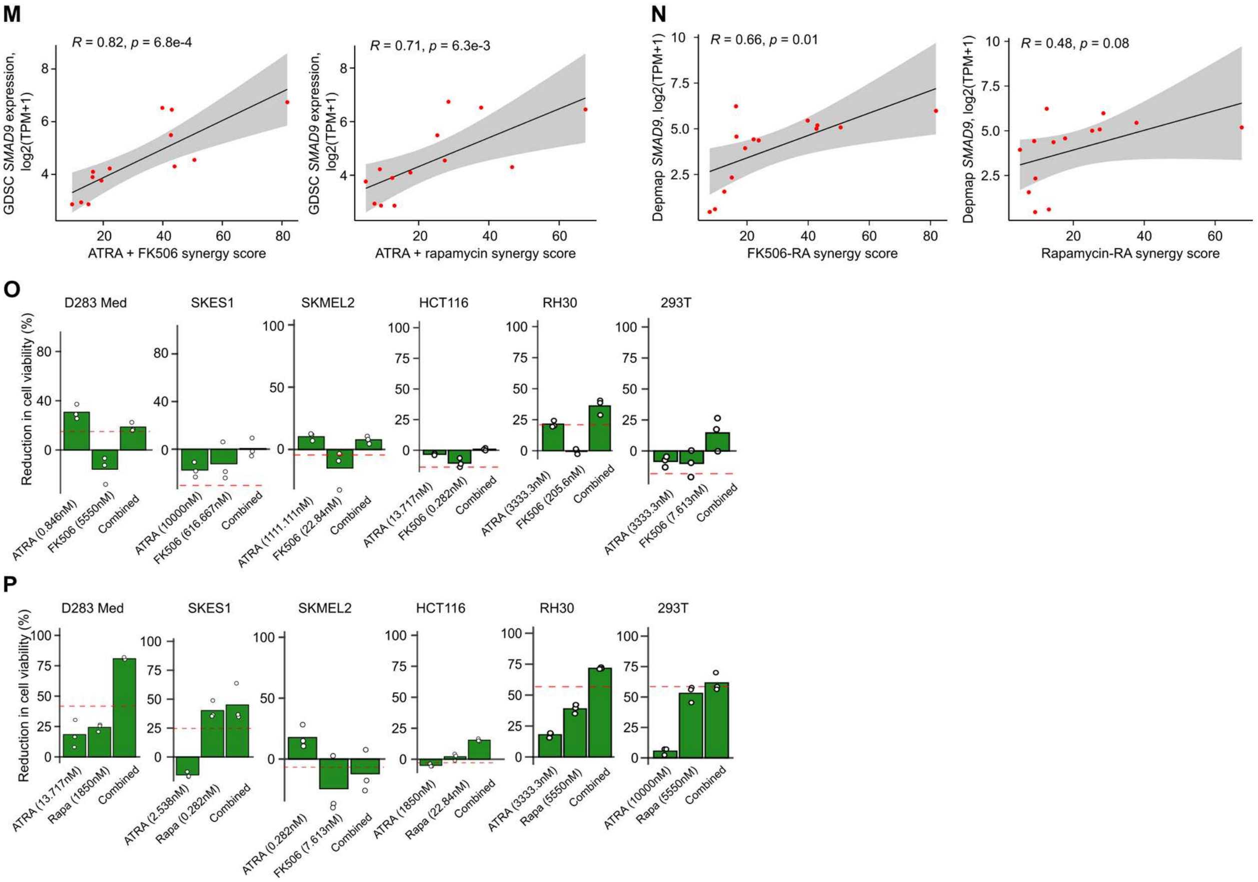


### Supplementary Figure 2

- **A.** *SMAD9* expression in all non-hematological cancer cell lines in the data from DepMap Expression Public 22Q4 (https://depmap.org/portal/gene/SMAD9?tab=characterization). Cancer types with ≤ 3 cell lines are not included.
- B. Kaplan-Meier curves showing overall survival in Cangelosi *et al.* (2020), grouped by *SMAD9* expression. For plotting purposes samples in the bottom 20% are designated as low expression. $n$ = 625 and $n$ = 156 samples in the low and high groups respectively. The "+" indicates censoring events. $P$-values calculated using a Cox proportional hazards model.
- C. Forest plot displaying the hazard ratio for *SMAD9* expression, along with other factors such as *MYCN* status, age, and INSS tumor stage. These hazard ratios were derived from a multivariate Cox proportional hazard model that jointly considers the association between overall survival and each of these factors.
- D. Scatter plot of *SMAD1* RNA expression from GDSC (left) and Depmap (right) plotted against RA IC50 values. P-values were calculated with two-sided Pearson correlation tests.
- E. Same as (A) but showing *SMAD5* expression and IC50.
- F. Western blots showing SMAD9 protein levels in CHP-134 and TGW cell lines following CRISPR knockout. Control, negative control gRNA with no target in human genome; All, a mixture of 3 independent gRNAs targeting *SMAD9*; 2972, a single gRNA targeting *SMAD9*. GAPDH was used as loading control. Representative results of 3 independent replicates are shown.

G. Viability of CHP-134 cells following *SMAD9* knockout under treatment with RA for 3 days (left) or 6 days (right). Data represent the mean ± SD of 3 independent replicates.
H. Viability of TGW cells following SMAD9 knockout under treatment with RA for 6 days. Data represent the mean ± SD of 3 independent replicates.
I. Western blots showing SMAD1 protein levels in CHP-134 and TGW cell lines following CRISPR knockout. Control, negative control gRNA with no target in human genome; All, a mixture of 3 independent gRNAs targeting *SMAD1*; 7252 and 1172 are single gRNAs targeting *SMAD1*. GAPDH was used as a loading control. Representative results of 3 independent replicates are shown.
J. Viability of CHP-134 (left) and TGW (right) cells following *SMAD1* knockout under treatment with RA for 6 days. Data represent the mean ± SD of 3 independent replicates.
K. Neuroblastoma cell lines were screened with combination of ATRA and FK506 as shown in Figure 2F-G. Bar plots of RA and FK506 combinations that conferred the maximum synergy scores from synergy calculations are shown here. White dots represent three independent experiments corresponding to score maxima. Red lines represent the expected result based on additivity alone.
L. Same as (K) but showing RA and rapamycin combinations.
M. Scatter plot of SMAD9 RNA expression from GDSC plotted against highest synergy score of ATRA + FK506 combination (left) and RA + rapamycin combination (right) in the cell lines screened in our drug combination study shown in Fig. E. P-values were calculated with two-sided Pearson correlation tests.
N. Same as (M) but showing *SMAD9* expression from Depmap.
O. Same as (K) but in non-neuroblastoma cell lines.
P. Same as (K) but showing RA and rapamycin combinations in non-neuroblastoma cell lines.

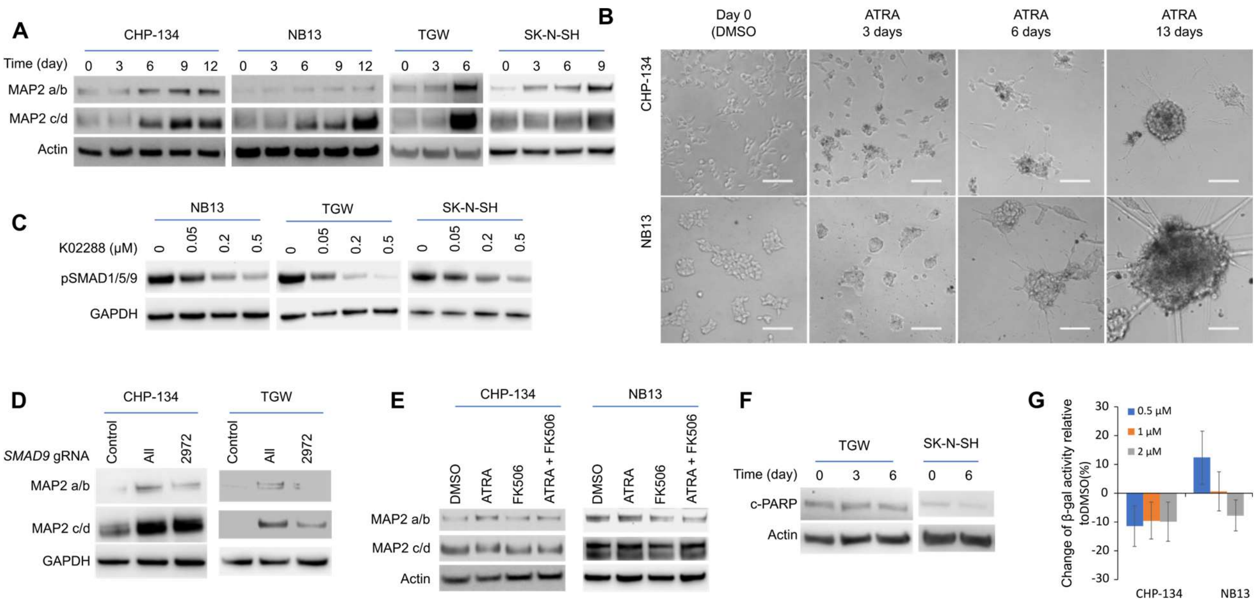


### Supplementary Figure 3

A. Western blots showing MAP2 expression in CHP-134, NB13, TGW, and SK-N-SH cell lines treated with 0.5 μM (CHP-134) or 2μM (NB13, TGW, SK-N-SH) of ATRA for the time indicated.
B. Images showing neurite growth in CHP-134 and NB13 cell lines after 6 days of RA treatment (0.5 μM for CHP-134 and 2 μM for NB13). Scale bar = 100 μm.
C. Western blots showing BMP inhibitor K02288 suppressed BMP signaling activity (pSMAD1/5/9 level) in NB13, TGW, and SK-N-SH cell lines. Cells were treated with K02288 with indicated concentrations for 3 days.
D. Western blots for MAP2 expression in CHP-134 and TGW cell lines after SMAD9 CRISPR knockout. Control, negative control gRNA with no target in human genome; All, a mixture of 3 independent gRNAs targeting SMAD9; 2972, a single gRNA targeting SMAD9.
E. Western blots for MAP2 expression in CHP-134 and NB13 cell lines treated with indicated compounds for 3 days (2 μM RA and 0.1 μM FK506 for CHP-134, 0.05 μM RA and 5 μM FK506 for NB13).
F. Western blots for c-PARP expression in TGW and SK-N-SH cell lines treated with RA (2 μM for TGW and 5 μM for SK-N-SH) for the time indicated.
G. Senescence levels quantified by intensity of fluorescence in CHP-134 and NB13 cell lines treated with RA for 6 days. Data represent the mean ± SD of 3 independent replicates.

For panel A and C-F, β-Actin and GAPDH were used as loading control for western blotting. Representative results of 3 independent replicates are shown.

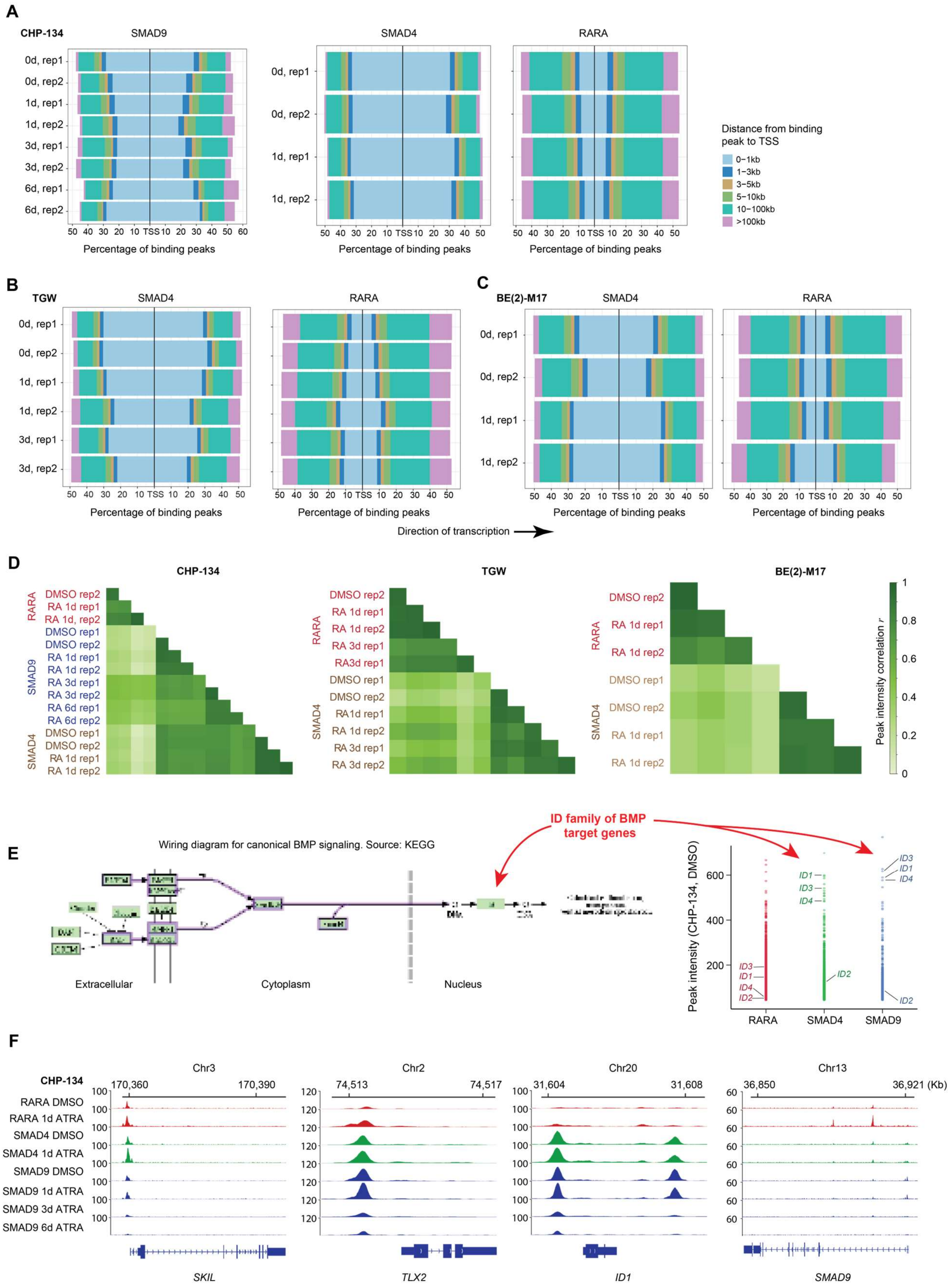

A
CHP-134
SMAD9
SMAD4
RARA
0d, rep1
0d, rep2
1d, rep1
1d, rep2
3d, rep1
3d, rep2
6d, rep1
6d, rep2
Percentage of binding peaks
Distance from binding peak to TSS
0–1kb
1–3kb
3–5kb
5–10kb
10–100kb
>100kb
B
TGW
C
BE(2)-M17
Direction of transcription
D
CHP-134
TGW
BE(2)-M17
Peak intensity correlation r
E
Wiring diagram for canonical BMP signaling. Source: KEGG
ID family of BMP target genes
Extracellular
Cytoplasm
Nucleus
Peak intensity (CHP-134, DMSO)
ID1
ID2
ID3
ID4
RARA
SMAD4
SMAD9
F
Chr3
Chr2
Chr20
Chr13
RARA DMSO
RARA 1d ATRA
SMAD4 DMSO
SMAD4 1d ATRA
SMAD9 DMSO
SMAD9 1d ATRA
SMAD9 3d ATRA
SMAD9 6d ATRA
SKIL
TLX2
ID1
SMAD9

**Supplementary Figure 4: Global summary/QC plots for the ChIP-seq data collected.**

A. The percentage of binding peaks (x axis) at various distances from transcription start sites (TSS; colors). Data shown for SMAD9 (left), SMAD4 (middle), and RARA (right) ChIP-seq profiles in CHP-134. Each row represents a sample, with the sample name indicating the duration of 2 μM ATRA treatment (0, 1, 3, or 6 days) and the replicate number (rep 1 or 2) number. RARA bindings sites are more dispersed across the genome, compared to SMAD profiles.
B. Like A but for TGW. Cells were treated with DMSO or 2 μM ATRA for one day or 0.5 μM ATRA for three days.
C. Like A but for BE(2)-M17. Cells were treated with DMSO or 2 μM ATRA for one day.
D. Hierarchical clustering of Pearson's correlations of shared-peak binding intensities between each pair of ChIP-seq samples for CHP-134 (left), TGW (middle) and BE(2)-M17 (right). The color indicates the factor profiled. In all cases, the strongest clustering is by technical replicate, followed by ChIP-mark and ATRA treatment, indicating the reproducibility/validity of our ChIP-seq data.
E. Stripchart (right panel) showing the genes bound by RARA, SMAD4, and SMAD9 in DMSO treated CHP-134 cells (x axis). The y axis indicates the peak binding intensity. As expected, the ID family of genes (the canonical targets of BMP signaling; left panel) are strongly bound by SMAD4 and SMAD9 in our data, indicating the data are reliable.
F. Selected genome browser tracks displaying the binding of RARA, SMAD4, and SMAD9 on four BMP target genes (*SKIL*, *TLX2*, *ID1*, and *SMAD9*) under DMSO or ATRA treatment in CHP-134, illustrating co-binding of these factors at many sites. SMAD9 binding is also lost over the treatment course, consistent with loss of SMAD9 mRNA expression in our RNA-seq data. Numbers on y-axis indicate the maximum normalized peak intensity.

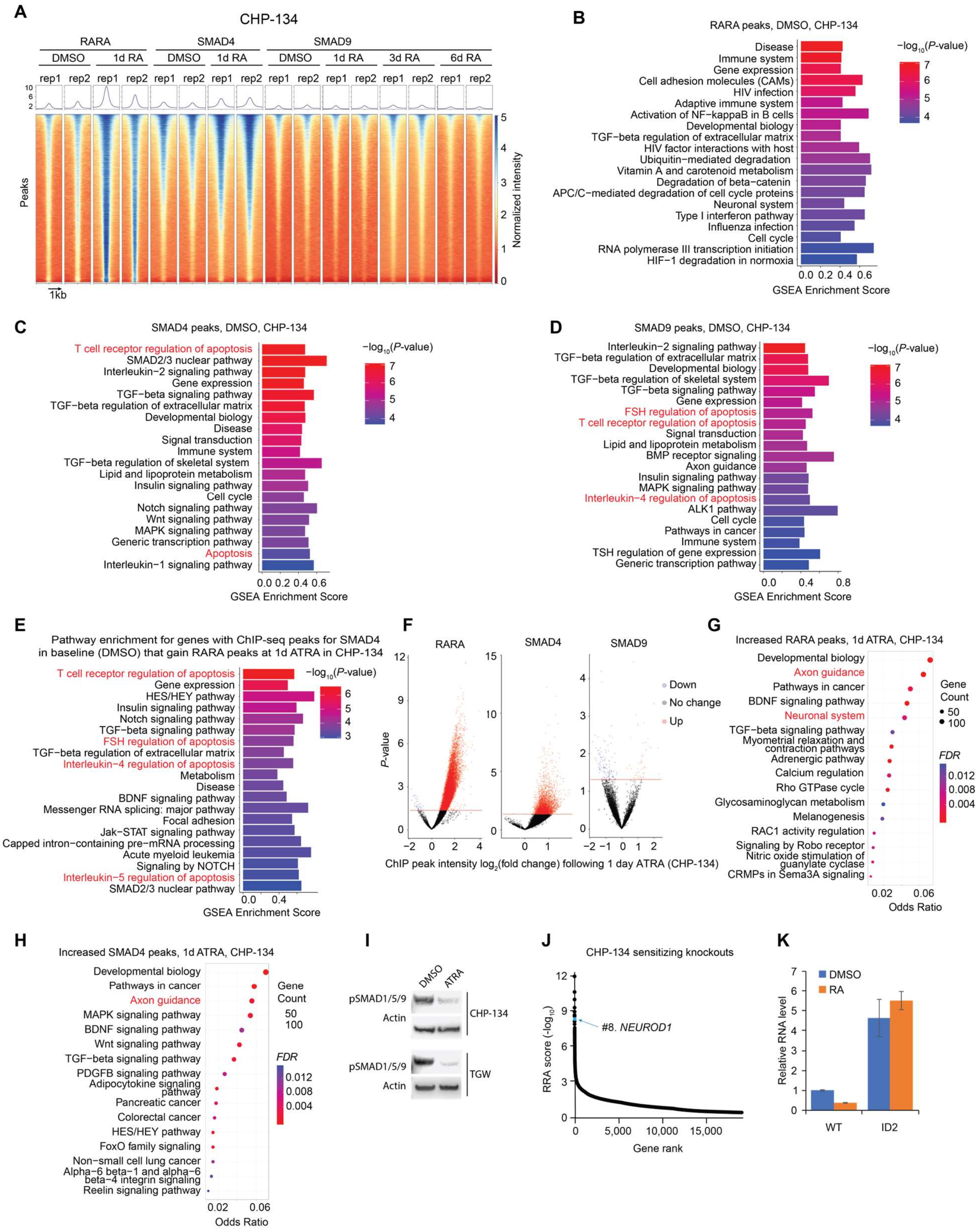


**Supplementary Figure 5: Additional analyses of ChIP-seq binding profiles in baseline and ATRA treated CHP-134 cells.**

A. Like Fig. 5A but SMAD9 ChIP-seq data from 3-day and 6-day treated (2 μM ATRA) samples are included. Genome-wide ChIP-seq binding intensity of RARA, SMAD4, and SMAD9 in CHP-134 cells following treatment with DMSO control or 2 μM ATRA for one day, three days, or six days are shown. Each row represents a genomic region, sorted by average binding intensity across all transcription factors. The color scale represents binding intensity, scale between 0 and 5 (see Methods). The binding profile (rows) of each sample was centered on a binding peak and shows 1kb upstream and downstream sequences from the peak.

B. Bar plot of GSEA enrichment scores (x axis) for BioPlanet pathways in DMSO treated CHP-134 cells, ranking genes based on the intensity of their RARA ChIP-seq peaks. The pathways are ordered by their *P*-value (color scale).

C. Like (B), but for SMAD4.

D. Like (B), but for SMAD9.

E. Bar plot (left panel) shows top GSEA enriched BioPlanet pathways for SMAD4 bound genes that gain RARA peaks following 2 μM ATRA treatment for 1 day (compared to DMSO) in CHP-134 cells.

F. Volcano plot of the changes in peak intensity of RARA, SMAD4, and SMAD9 in CHP-134 cells following 2 μM ATRA exposure for 1 day (compared to DMSO). Peaks with a $\log_2$ fold change (x axis) of >1 or < -1 and an adjusted *P*-value < 0.05 are highlighted as significant (colors).

G. Dot plot of the top enriched pathways for genes whose RARA peak intensity increased with 2 μM ATRA treatment for 1 day (compared to DMSO). *P*-values and odds ratios (x axis) were calculated by hypergeometric test.

H. Like (G), except for genes whose SMAD4 peak intensity increased with RA treatment.

I. Western blots showing expression of pSMAD1/5/9 in CHP-134 and TGW cells treated with 2 μM ATRA for 3 days.

J. Waterfall plot showing top hits involved in BMP signaling, ranked by negative RRA score (a higher value on y-axis indicates the gene knockout is more likely to increase CHP-134 sensitivity to ATRA). *NEUROD1* is highlighted.

K. *ID2* expression in wild-type (WT) and *ID2*-overexpressed (*ID2*) CHP-134 cells following DMSO or 2 μM RA for six days. Data represents the mean ± SD of three independent replicates.

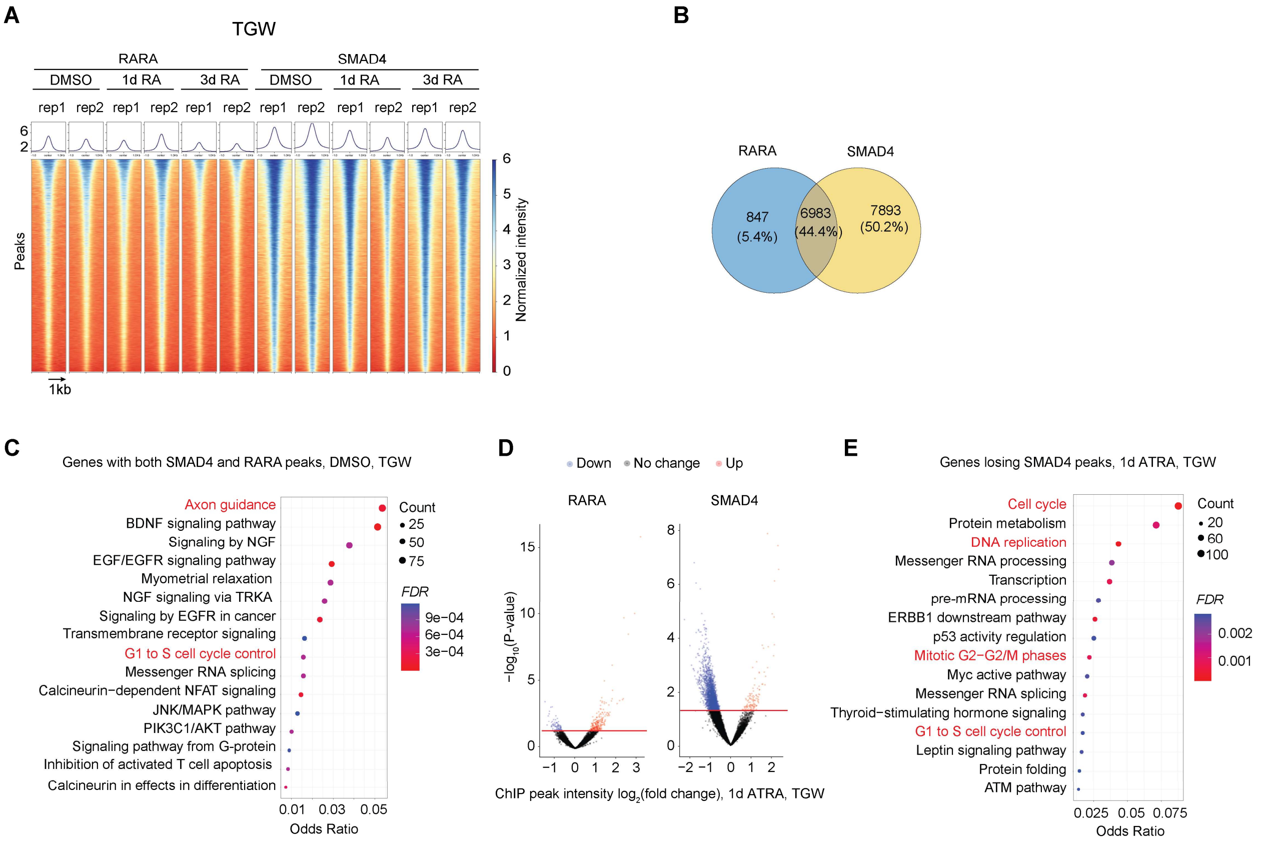


**Supplementary Figure 6: Additional analyses of ChIP-seq binding profiles in baseline and ATRA treated TGW cells.**

A. Heatmap showing RARA and SMAD4 binding ChIP-seq profiles in TGW cells. The cells were treated with DMSO, 2 μM ATRA for one day, or 0.5 μM ATRA for three days. Two independent replicates are shown.
B. Venn diagram showing the number and proportion of overlapping binding peaks between RARA and SMAD4 in TGW.
C. Dot plot showing the most enriched pathways for genes bound by both SMAD4 and RARA in DMSO treated TGW cells. *P*-values and odds ratios (x axis) were calculated by hypergeometric test.
D. Volcano plot showing intensity change of RARA and SMAD4 binding peaks in TGW cells in response to 2 μM ATRA exposure for 1 day (compared to DMSO). Peaks with a $\log_2$ fold change > 1 or < -1 and an *FDR* <0.05 were considered significant (colors).
E. Dot plot showing the most enriched pathways for genes whose SMAD4 peak intensity decreased significantly with 2 μM ATRA treatment for 1 day. *P*-values and odds ratios (x axis) were calculated by hypergeometric test.

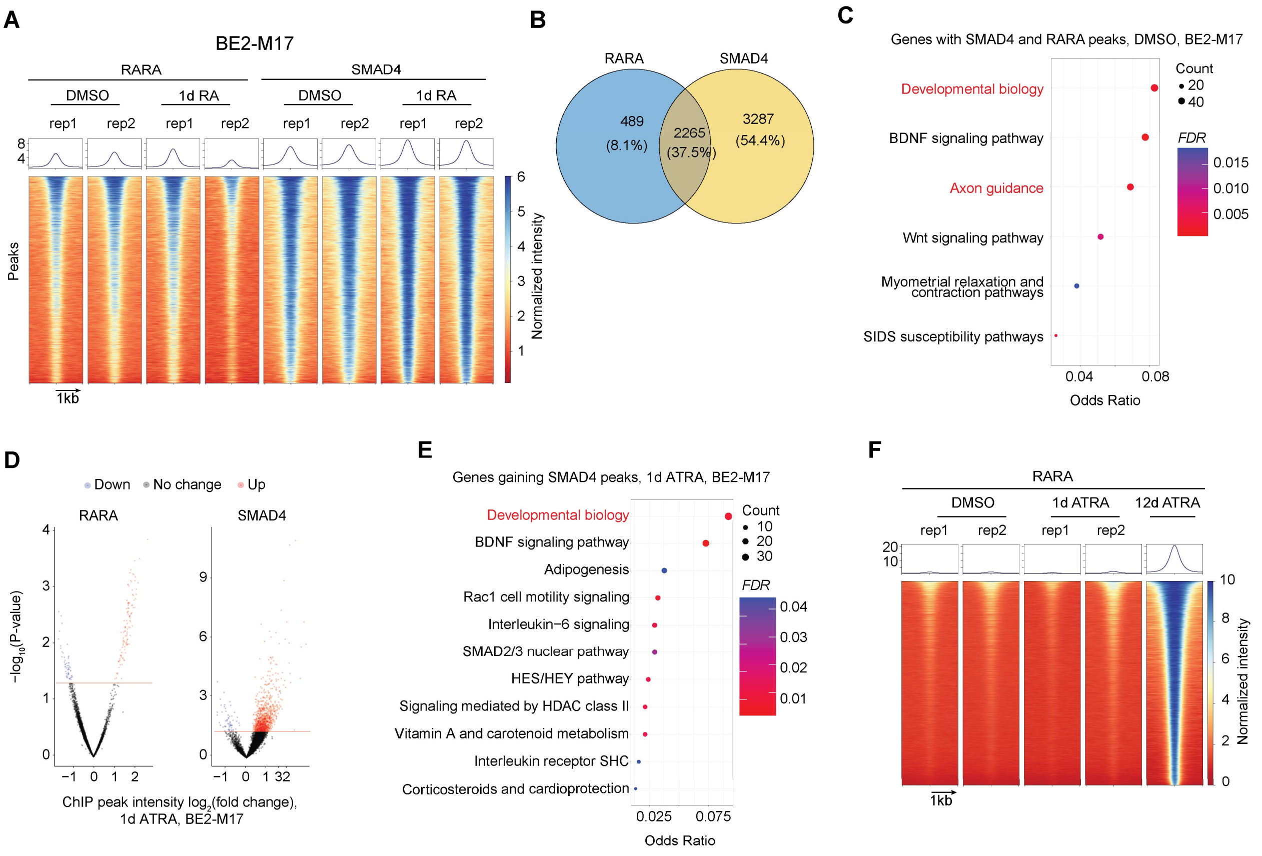


**Supplementary Figure 7: Additional analyses of ChIP-seq binding profiles in baseline and ATRA treated BE(2)-M17 cells.**

A. Heatmap showing RARA and SMAD4 binding profiles for all data collected in BE(2)-M17 cells.
B. Venn diagram showing the number and proportion of overlapping binding peaks between RARA and SMAD4 in BE(2)-M17.
C. Dot plot showing the most enriched pathways for genes bound by both SMAD4 and RARA in baseline DMSO treated BE(2)-M17 cells. *P*-values and odds ratios (x axis) calculated by hypergeometric test.
D. Volcano plot showing changes in peak intensity of RARA and SMAD4 in BE(2)-M17 cells in response to 2 μM ATRA exposure for 1 day. Peaks with a $\log_2$ fold change > 1 or < -1 and an FDR <0.05 were considered significant (colors).
E. Volcano plot showing changes in peak intensity of RARA and SMAD4 in BE(2)-M17 cells in response to 2 μM ATRA exposure for 1 day. Peaks with a $\log_2$ fold change > 1 or < -1 and an *FDR* <0.05 were considered significant (colors).
F. Heatmap displaying the genome-wide ChIP-seq binding intensity of RARA in BE(2)-M17 following treatment with DMSO control or 2 μM ATRA for 1 d, in comparison to the RARA binding profile in BE2C cell line treated with 5 μM ATRA for 12 days, where these cells eventually gain RARA peaks.

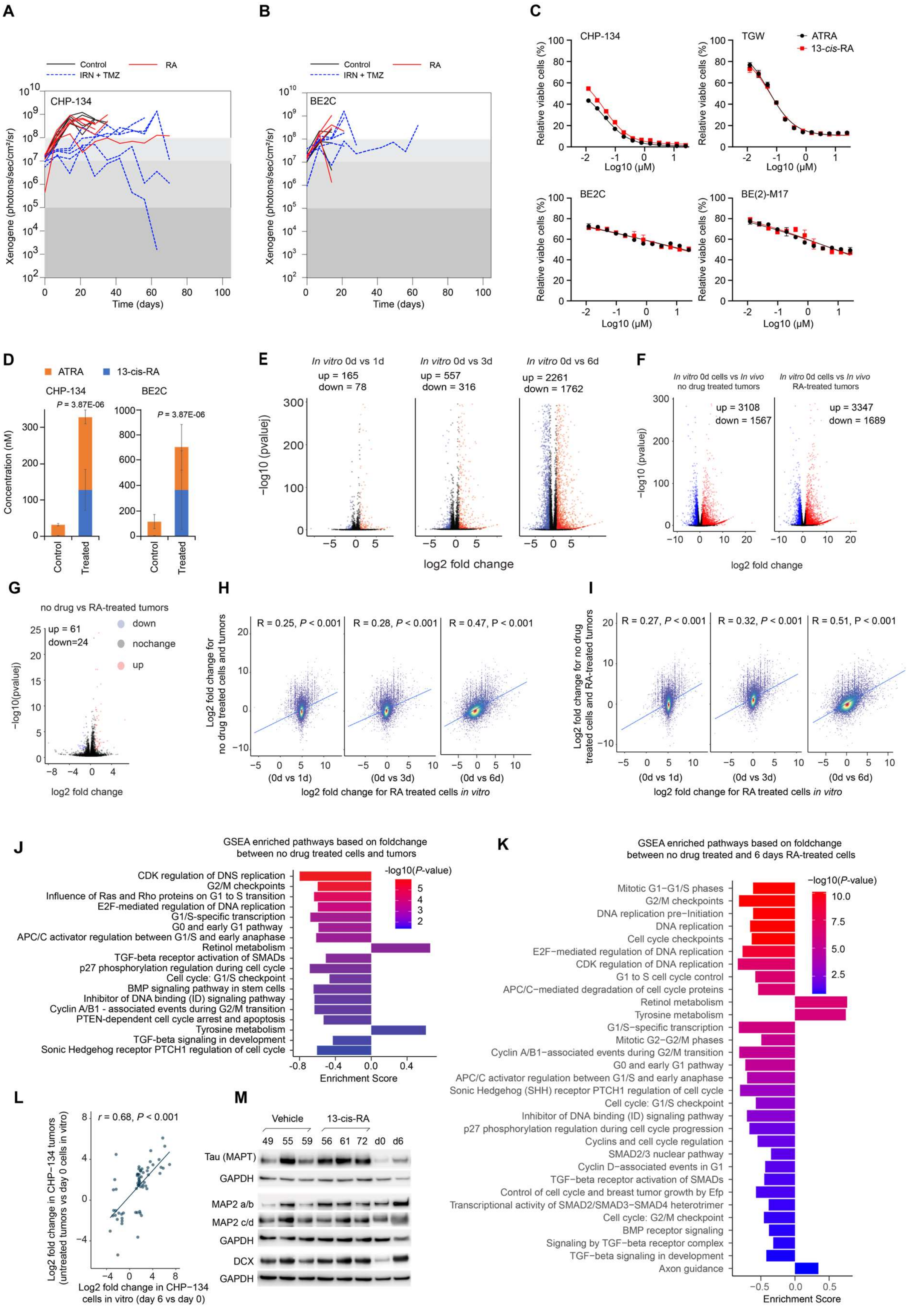
A
B
C
Control
RA
IRN + TMZ
CHP-134
BE2C
Xenogene (photons/sec/cm²/sr)
Time (days)
CHP-134
TGW
BE2C
BE(2)-M17
ATRA
13-cis-RA
Relative viable cells (%)
Log10 (μM)
D
ATRA
13-cis-RA
CHP-134
BE2C
P = 3.87E-06
Concentration (nM)
Control
Treated
E
In vitro 0d vs 1d
up = 165
down = 78
In vitro 0d vs 3d
up = 557
down = 316
In vitro 0d vs 6d
up = 2261
down = 1762
−log10 (pvaluej)
log2 fold change
F
In vitro 0d cells vs In vivo no drug treated tumors
In vitro 0d cells vs In vivo RA-treated tumors
up = 3108
down = 1567
up = 3347
down = 1689
−log10 (pvaluej)
log2 fold change
G
no drug vs RA-treated tumors
up = 61
down=24
down
nochange
up
−log10(pvalue)
log2 fold change
H
R = 0.25, P < 0.001
R = 0.28, P < 0.001
R = 0.47, P < 0.001
Log2 fold change for no drug treated cells and tumors
(0d vs 1d)
(0d vs 3d)
(0d vs 6d)
log2 fold change for RA treated cells in vitro
I
R = 0.27, P < 0.001
R = 0.32, P < 0.001
R = 0.51, P < 0.001
Log2 fold change for no drug treated cells and RA-treated tumors
(0d vs 1d)
(0d vs 3d)
(0d vs 6d)
log2 fold change for RA treated cells in vitro
J
GSEA enriched pathways based on foldchange between no drug treated cells and tumors
CDK regulation of DNS replication
G2/M checkpoints
Influence of Ras and Rho proteins on G1 to S transition
E2F-mediated regulation of DNA replication
G1/S-specific transcription
G0 and early G1 pathway
APC/C activator regulation between G1/S and early anaphase
Retinol metabolism
TGF-beta receptor activation of SMADs
p27 phosphorylation regulation during cell cycle
Cell cycle: G1/S checkpoint
BMP signaling pathway in stem cells
Inhibitor of DNA binding (ID) signaling pathway
Cyclin A/B1 - associated events during G2/M transition
PTEN-dependent cell cycle arrest and apoptosis
Tyrosine metabolism
TGF-beta signaling in development
Sonic Hedgehog receptor PTCH1 regulation of cell cycle
-log10(P-value)
Enrichment Score
K
GSEA enriched pathways based on foldchange between no drug treated and 6 days RA-treated cells
Mitotic G1−G1/S phases
G2/M checkpoints
DNA replication pre−Initiation
DNA replication
Cell cycle checkpoints
E2F−mediated regulation of DNA replication
CDK regulation of DNA replication
G1 to S cell cycle control
APC/C−mediated degradation of cell cycle proteins
Retinol metabolism
Tyrosine metabolism
G1/S−specific transcription
Mitotic G2−G2/M phases
Cyclin A/B1−associated events during G2/M transition
G0 and early G1 pathway
APC/C activator regulation between G1/S and early anaphase
Sonic Hedgehog (SHH) receptor PTCH1 regulation of cell cycle
Cell cycle: G1/S checkpoint
Inhibitor of DNA binding (ID) signaling pathway
p27 phosphorylation regulation during cell cycle progression
Cyclins and cell cycle regulation
SMAD2/3 nuclear pathway
Cyclin D−associated events in G1
TGF−beta receptor activation of SMADs
Control of cell cycle and breast tumor growth by Efp
Transcriptional activity of SMAD2/SMAD3−SMAD4 heterotrimer
Cell cycle: G2/M checkpoint
BMP receptor signaling
Signaling by TGF−beta receptor complex
TGF−beta signaling in development
Axon guidance
-log10(P-value)
Enrichment Score
L
r = 0.68, P < 0.001
Log2 fold change in CHP−134 tumors (untreated tumors vs day 0 cells in vitro)
Log2 fold change in CHP−134 cells in vitro (day 6 vs day 0)
M
Vehicle
13-cis-RA
49 55 59 56 61 72 d0 d6
Tau (MAPT)
GAPDH
MAP2 a/b
MAP2 c/d
GAPDH
DCX
GAPDH

**Supplementary Figure 8**

A. Mice were implanted with CHP-134 cells. Figure shows the photon emissions of the resulting tumors (as measured by ultrasound). Greater photon emissions indicate greater tumor volume. Curves are colored by the treatment group.
B. Same as in (A), but the tumors were generated by implanting mice with BE2C cells.
C. Viability of cell lines (CHP-134, TGW, BE2C, BE(2)-M17) treated for six days with *all-trans*-retinoic acid (ATRA) or 13-cis-retinoic acid (13-*cis*-RA). Data represents the mean ± SD of three independent experiments.
D. Concentrations of ATRA and 13-*cis*-RA in vehicle (Control) and 13-*cis*-RA-treated (Treated) tumor tissues. Tumors were collected within two weeks of the last treatment. The number of tumors in group are as follows: four in CHP-134 Control; three in CHP-134 Treated; five in BE2C Control; and four in BE2C Treated. P-values were calculated using two-tailed t tests.
E. Volcano plot of the differentially expressed genes in CHP-134 cells treated with RA for 1, 3, and 6 days versus control CHP-134 cells (0 days). Cells were cultured *in vitro.* Genes were considered differentially expressed if they exhibited a log2 fold change greater than 1 or less than -1 as well as an adjusted p-value of less than 0.05.
F. Left: volcano plot of the differentially expressed genes in untreated CHP-134 tumors vs untreated CHP-134 cells cultured *in vitro*. Right: volcano plot of the differentially expressed genes in CHP-134 tumors treated with RA vs untreated CHP-134 cells cultured in *vitro.*
G. Volcano plot of the differentially expressed genes in RA-treated vs untreated (control) CHP-134 tumors.
H. Scatter plots comparing the gene expression fold change between untreated CHP-134 cells cultured *in vitro* and untreated CHP-134 tumors versus the gene expression fold change between untreated CHP-134 cells cultured *in vitro* and CHP-134 cells treated with RA for 1 day (left panel), 3 days (middle panel) and 6 days (right panel) *in vitro*. Pearson correlation coefficients and p-values are displayed on the plots.
I. Similar to (I), but the values on the y-axis represent the fold change between untreated CHP-134 cells cultured *in vitro* and RA-treated CHP-134 tumors.
J. Bar plot of the enriched pathways identified through GSEA analysis. In this analysis, genes were ranked by the fold change in their expression in untreated CHP-134 cells cultured *in vitro* versus untreated CHP-134 tumors.
K. Bar plot of the enriched pathways identified through GSEA analysis. In this analysis, genes were ranked by the fold change in their expression in untreated CHP-134 cells cultured *in vitro* versus CHP-134 cells treated with RA for six days *in vitro*.
L. For differentially expressed genes enriched in axon guidance pathway, as shown in Fig. 6E, their expression fold change between untreated CHP-134 cells cultured *in vitro* and untreated CHP-134 tumors *in vivo* are plotted against the expression fold change between untreated CHP-134 cells cultured *in vitro* and CHP-134 cells treated with RA for 6 days. Pearson correlation coefficients and *P*-values are displayed on the plots.
M. Western blots showing the expression of neuron markers MAPT, MAP2, and DCX in CHP-134 xenograft tumor tissues and the cell line cultured in vitro (treated with DMSO (d0) or 2 μM RA for 6 days (d6)). GAPDH was used loading control. The representative results of 3 independent replicates are shown.

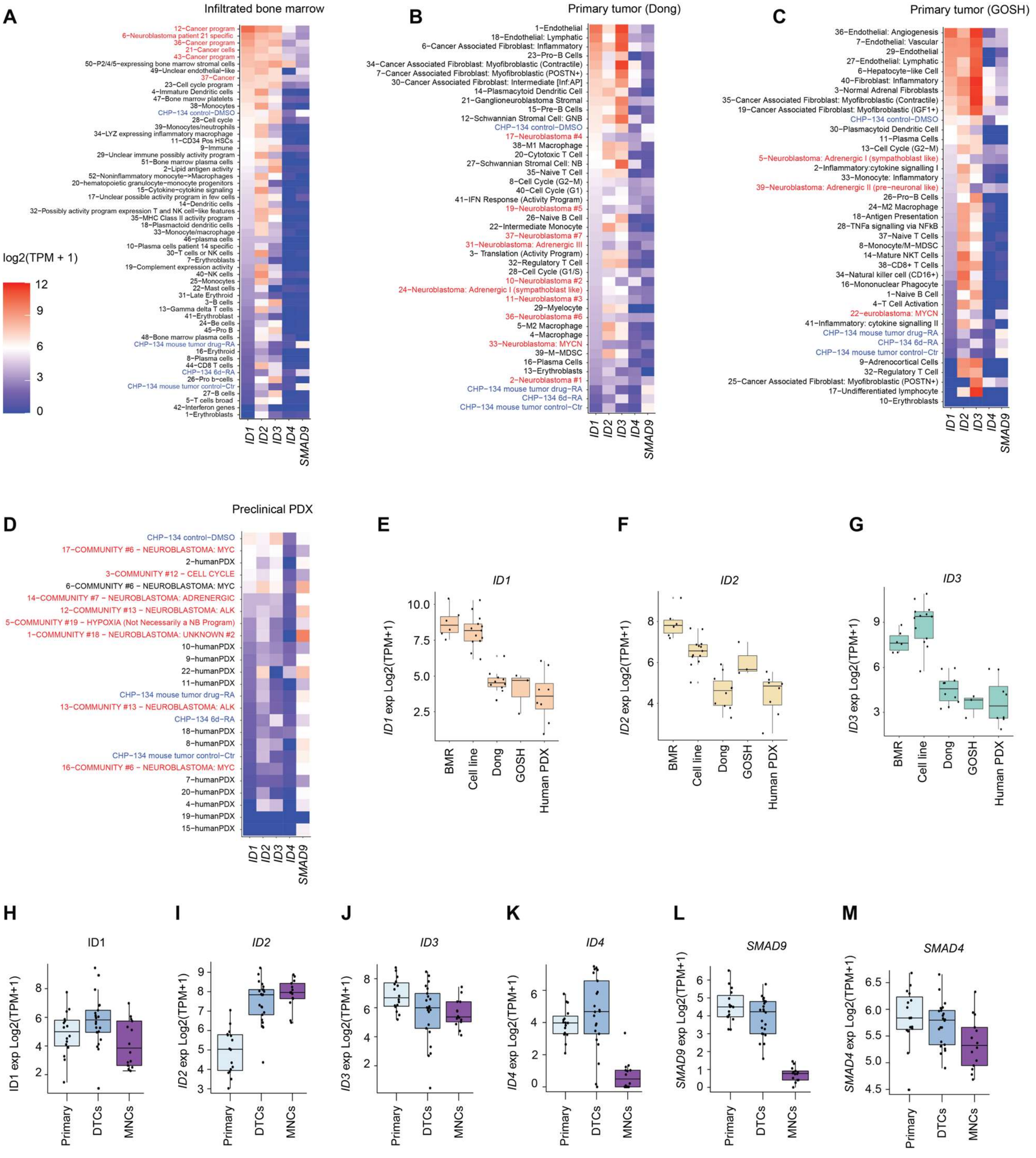


### Supplementary Figure 9

A. Heatmap showing the expression levels of *ID* genes and *SMAD9* in cells expression 52 gene expression programs (cell clusters categorized by gene expression profiles) identified from single-cell RNA-sequencing data of neuroblastoma-infiltrated bone marrow samples (data from Fetahu *et al*, PMID 37365178). Programs highlighted in red represent those expressed in neuroblastoma cancer cells, while those in black indicate tumor-associated normal cells. Blue represents CHP-134 cells and *in vivo* tumors, which have been integrated into the plot to serve as a reference for expression comparison with the bone marrow cancer cells.

B. Similar to (A) but for single-cell RNA-seq in primary neuroblastoma tumors (data from Dong et al, PMID 32946775).
C. Similar to (A) but for single-cell RNA-seq in another primary neuroblastoma dataset (data from Kildisuite *et al*, PMID 33547074).
D. Similar to (A) but for single-cell RNA-seq of a preclinical PDX dataset (data from Chapple *et al*.).
E. Boxplots showing the expression of *ID1* in cells expressing the cancer programs (programs highlight by red color in A-D) identified in single-cell RNA-seq from bone marrow (BMR), human primary neuroblastoma tumors (Dong and GOSH), PDXs model (Human PDX) and 13 neuroblastoma cell lines (Cell line).
F. Similar to (E) but for the expression of *ID2*.
G. Similar to (E) but for the expression of *ID3*.
H. Boxplots (H-M) showing $log_2$(TPM + 1) normalized bulk RNA-seq expression (y axis) of key BMP pathway genes *ID1-4* (H-K), *SMAD9* (L) and *SMAD4* (M) in primary human neuroblastoma tumors and bone marrow metastatic sites from the same patients. Boxes represent bulk primary tumors (‘Primary’; light blue), disseminated tumors cells (‘DTCs’; dark blue) in the bone marrow enriched using an anti-GD2 antibody, and the remaining mononuclear cells (‘MNCs’; purple) following depletion of GD2 expressing cells. Data were obtained Rifatbegovic *et al*. (PMID 28921546).

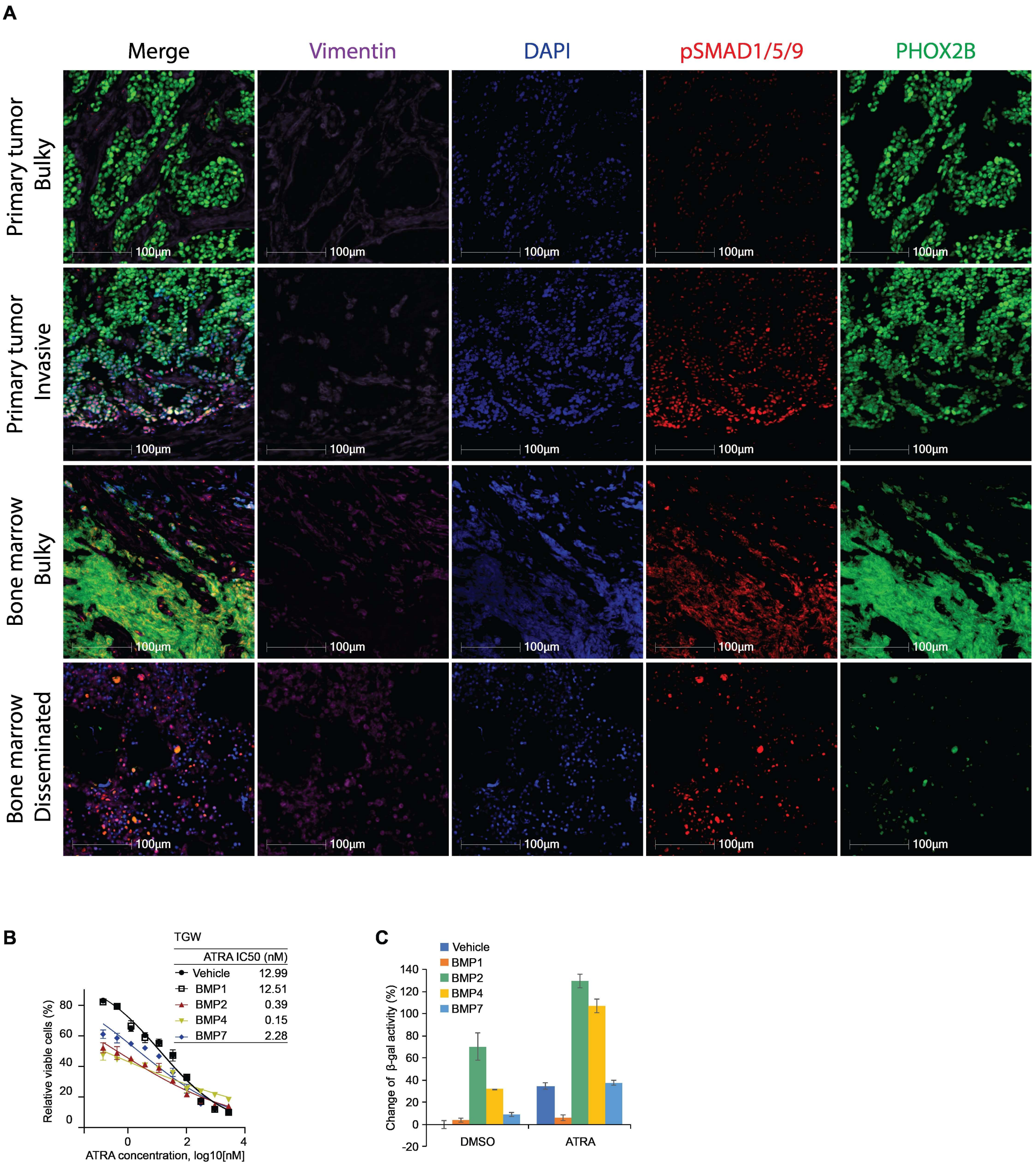


### Supplementary Figure 10

A. Related to Fig. 6H. Immunofluorescent images from Patient #3 showing Vimentin, DAPI, pSMAD1/5/9, and PHOX2B staining. The cells co-stained with pSMAD1/5/9 and PHOX2B appear yellow or orange in the merged images.
B. Viability of TGW cells treated with ATRA for 6 days in the presence of vehicle control or 10 ng/ml BMP recombinant proteins. Cell viability was measured with CellTiter-Glo.

ATRA IC50 for TGW (right) in the presence of each BMP was calculated with GraphPad Prism 9. Data represent the mean ± SD of 3 independent replicates.

C. Senescence levels quantified by intensity of fluorescence in TGW cells treated with DMSO and 0.03 μM ATRA for 6 days in the presence of vehicle control or 10 ng/ml BMP recombinant proteins. Data represent the mean ± SD of 3 independent replicates.
